# Comparing instance segmentation methods for analyzing clonal growth of single cells in microfluidic chips

**DOI:** 10.1101/2020.12.31.424955

**Authors:** Elliott D. SoRelle, Scott White, Benjamin B. Yellen, Kris C. Wood, Micah A. Luftig, Cliburn Chan

## Abstract

Appropriately tailored segmentation techniques can extract detailed quantitative information from biological image datasets to characterize and better understand sample distributions. Practically, high-resolution characterization of biological samples such as cell populations can provide insights into the sources of variance in biomarker expression, drug resistance, and other phenotypic aspects, but it is still unclear what is the best method for extracting this information from large image-based datasets. We present a software pipeline and comparison of multiple image segmentation methods to extract single-cell morphological and fluorescence quantitation from time lapse images of clonal growth rates using a recently reported microfluidic system. The inputs in all pipelines consist of thousands of unprocessed images and the outputs are the detection of cell counts, chamber identifiers, and individual morphological properties of each clone over time detected through multi-channel fluorescence and bright field imaging. Our conclusion is that unsupervised learning methods for cell segmentation substantially outperform supervised statistical methods with respect to accuracy and have key advantages including individual cell instance detection and flexibility through model training. We expect this system and software to have broad utility for researchers interested in high-throughput single-cell biology.

## Introduction

Imaging science has been foundational to biological investigation for centuries,^1,2^ enabling key discoveries from the scale of single molecules^3,4^ to whole organisms.^5-7^ Methodological advances in image acquisition and analysis have served as cornerstones for everyday benchtop assays and breakthrough technologies including high-throughput screening and sequencing platforms.^8,9^ Frequently, imaging approaches are used to apply binary classifications (signal versus background) to biological events (e.g., presence of a biomolecule of interest, labeled nucleotide incorporation) and count or otherwise quantify occurrences of biological phenomena. Less frequently, image analysis is deployed for extraction and high-dimensional characterization of features within biological samples, such as the size or shape of individual cells and their spatial context relative to other cells in a culture environment or tissue.^10-13^ Analytical methods that can provide multidimensional feature quantification are increasingly valuable tools for discovery. The high density of images in combination with machine learning and image segmentation models that are trained using well-annotated training sets can together be used to reveal insights that not only quantify intrinsic biological heterogeneity but also reveal deeper relationships that are not readily apparent. For example, the intra-sample diversity with respect to gene expression, drug resistance, and other phenomena can have substantial functional implications for the biological responses to perturbations.^14-17^

Cell populations (both primary and transformed lines) are one class of biological systems that could benefit greatly from granular, multiparametric characterization. Such populations can comprise individual cells with highly variant phenotypes that fluctuate over time, or rare phenotypes that can be identified in a background of more abundant phenotypes.^18-20^ Whereas technical capabilities have been developed to capture single-timepoint genetic and transcriptomic profiles of individual cells in a sample,^21-23^ methods for characterizing single-cell phenotypes in a high-throughput, time-resolved fashion have lagged because of the need to maintain and repeatedly measure the same cell over time. To address this technical gap, Celldom™ recently developed and reported a microfluidic platform that achieves highly parallel single-cell culture.^24^ By trapping individual cells deterministically, this system can be used in conjunction with a laboratory microscope to monitor growth dynamics of thousands of individual cells within a population.

In this work, we developed a software package and conducted a systematic comparison of methods to segment and count cells in images collected using the Celldom™ platform. In addition to creating an efficient pipeline for extracting clone-specific data, we systematically tested numerous image segmentation techniques to maximize cell classification performance. The package enables high-dimensional, time-resolved characterization of clonal dynamics, morphology, and biomarker expression when used in conjunction with standard fluorescence assays. We expect the fusion of these capabilities to expand the potential and versatility of single-cell biology experimentation with enhanced statistical power.

## Results

### Processing pipeline overview

The Celldom™ chip consists of a microfluidic network of >6000 culture chambers, each with a constricted opening that can physically trap and load exactly one cell per chamber under appropriate conditions.^24^ Each chamber is uniquely identified by a combination of numerical row and column addresses, which are etched into the chip at specific locations relative to each chamber. Additionally, the chip contains a repeating pattern of fiducial marks at fixed positions relative to each chamber and its corresponding row and column addresses. Every field of view (10x magnification) across the chip surface contains these components (**Figure 1A**), each of which can be automatically identified through image processing techniques. We developed an image processing pipeline to analyze Celldom™ image datasets to extract quantitative single-cell information for individual cultured clones, which included a systematic performance characterization of numerous methods for cell segmentation (**Figure 1B**). In the first step of the pipeline, the cross-shaped fiducial marks in each field of view are identified using template matching. An optional step for illumination normalization can be applied after fiducial detection. The centroids of these fiducials are used to apply image-wide small angle rotation correction and to define rectangular regions of interest (ROIs) containing the corresponding chamber, row, and column based on conserved translations in the microfluidic layout pattern (**Figure 1C**). Next, individual digits within the row and column ROIs are template-matched to a library of reference digits and recorded using a majority vote process for the optimal match. This method correctly identified chamber addresses >95% of the time for a typical experiment, which translates to >5,700 correctly indexed clonal growth chambers per chip. Addressed chambers are then analyzed using a segmentation method to produce masks of cell-containing regions (**Figure 1D**) and, in the case of the best-performing method, high-accuracy detection of individual cell instances within each chamber (**Figure 1E**), enabling intra- and inter-clonal single-cell phenotype comparisons.

**Figure 1.**
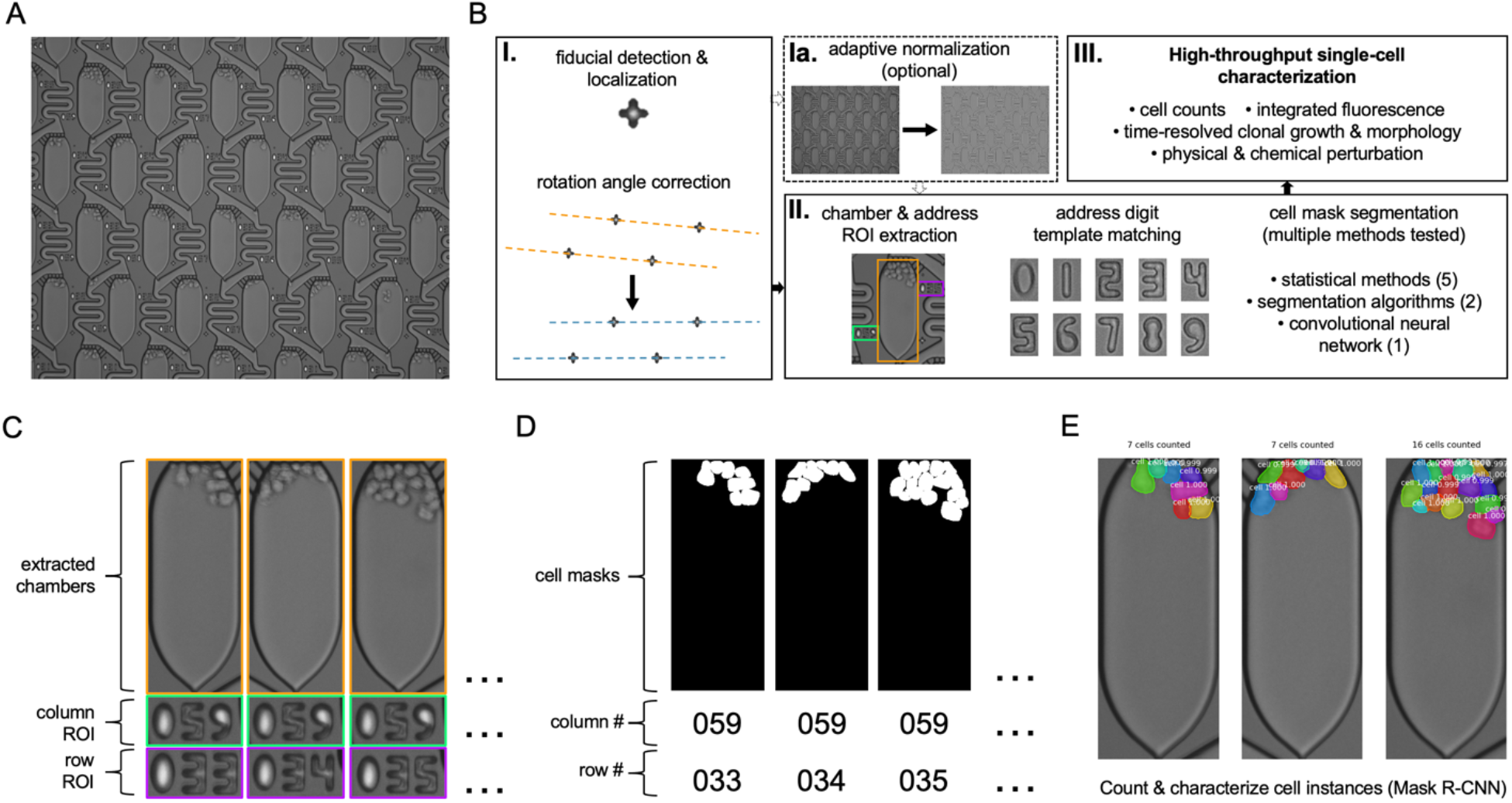
Image processing workflow for Celldom single-cell culture platform. (**A**) Example brightfield image of chamber cultures initiated from single B cells (LCL 461) on the Celldom microfluidic array. Chambers are connected in a microfluidic ladder network and are uniquely identified by etched row and column addresses. (**B**) Raw image-to-single-cell quantification processing pipeline for high-resolution cell biological studies. Images are pre-processed to extract etched fiducial marks and make fine image rotation corrections based on calculated fiducial row angles (box I). Optionally, light variation across the field of view can be reduced through local adaptive normalization methods at the expense of dynamic range (box Ia). After pre-processing corrections, culture chamber and corresponding address ROIs are extracted based on fixed positions relative to fiducial marks. Individual digits in the row and column ROIs are template-matched to a library of reference digits to read chamber addresses, and a given method for cell segmentation is applied to identify cells within the chamber ROI (box II). Results of chamber-indexed cell segmentation can be applied to collect high-dimensional data on individual cells to suit specific experimental setups (box III). (**C**) Representative examples of culture chamber, column, and row ROIs extracted after fiducial identification and pre-processing. (**D**) Corresponding chamber-indexed cell segmentation masks for the chambers shown in C. Masks in this example depict composites of individual cell instance identification results from the implementation of a trained model using mask-rcnn. (**E**) Color-coded mask-rcnn detection of individual cell instances, enabling highly parallel accurate counting and morphological characterization of cells within clonal populations.

### Method development and selection

Because the quality of image segmentation can have profound effects on the biological interpretation of an experiment,^25^ we sought quantitative characterization and comparison of multiple methods for identifying cells in extracted chambers (**Figure 2**). To this end, we evaluated the performance of five methods based on image statistics (Difference of Gaussians,^26^ Standard Deviation, Entropy,^27^ Inverse Canny Edge,^28^ and Rolling 2D Standard Deviation); two established segmentation algorithms (Chan-Vese^29^ and Felzenszwalb^30^); and one machine learning approach (Mask R-CNN^31^) to identify cells (**Figure 2A**).

**Figure 2.**
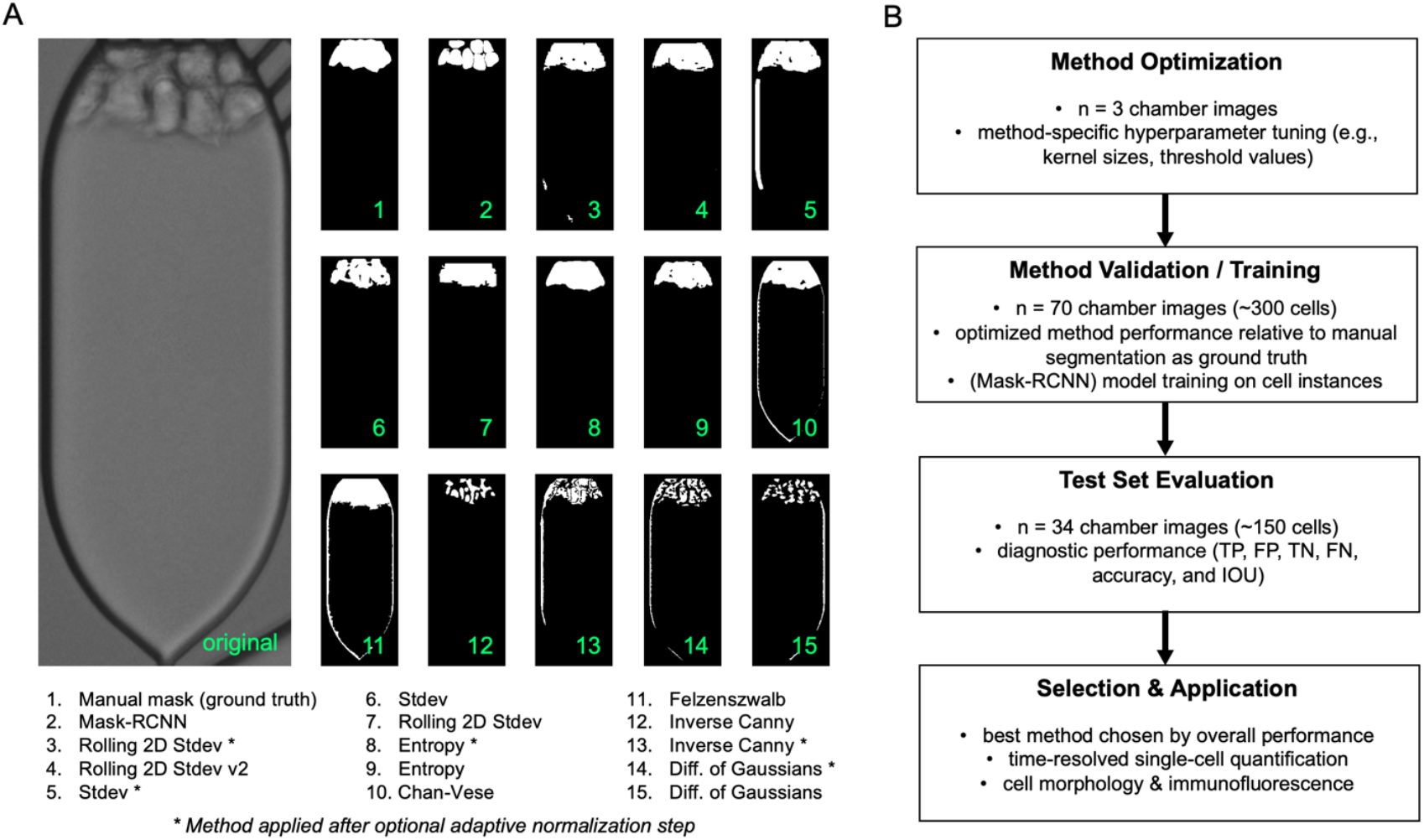
Comparative study of methods for cell segmentation in culture chambers. (**A**) Cell segmentation masks were generated using various methods and evaluated relative to manually segmented ground truth masks. Five statistical methods for cell segmentation (Rolling 2D Stdev, Stdev, Entropy, Inverse Canny, and Difference of Gaussians) were developed and applied to images with (*) and without adaptive normalization. Additionally, two existing segmentation algorithms (Chan-Vese and Felzenszwalb) were tested. A machine learning approach using mask-rcnn and a trained cell detection model was also evaluated. (**B**) After initial development, each of the five statistical methods and the two segmentation algorithms were independently optimized for extracted chamber ROIs through hyperparameter tuning. Optimized performance of each method was confirmed using a validation set of 70 chamber images. In the case of the mask-rcnn method, individual cells within these 70 images were used for model training. The trained model was initiated from mask-rcnn’s coco dataset model and was generated using 30 training epochs. After validation and training, each method was applied to a test set of 34 chamber images to generate reported performance metrics. The best method based on diagnostic criteria was selected and implemented for subsequent quantitative studies of single-cell culture experiments.

We designed a study to optimize and quantitatively assess the performance of each of the selected cell segmentation methods (**Figure 2B**). After tuning method-specific hyperparameters (see **Materials and Methods**) on a small set of preliminary images (n = 3), 104 images of individual extracted chambers containing transformed human B cells (a lymphoblastoid cell line, LCL 461) were divided into validation/training (n = 70) and test (n = 34) sets. We manually segmented reference masks of cell-containing area in all images within each set to use as a ground truth classification standard. Mask-RCNN requires a trained model for classification, so we used individual cells segmented from the 70 validation set images for training (see **Materials and Methods**). After validating and optimizing method performance, we analyzed the test set images to generate quantitative performance evaluation for each segmentation method. The best method based on overall performance was implemented in the processing pipeline.

### Comparative performance evaluation

Based on hyperparameter tuning, methods were developed to favor specificity over sensitivity, which is generally reflected in the set-wide distributions of pixel-wise classification rates relative to ground truth reference masks (**Figure 3A**). Most of the statistical segmentation methods outperformed the two established segmentation algorithms (except for the Inverse Canny Edge and Difference of Gaussians methods). However, the Mask R-CNN model implementation was superior to all other methods with respect to each evaluated metric (**Figure 3A, B**). Measurements of intersection over union (IOU) of method classification results relative to reference masks indicate that Mask-RCNN and two versions of the Rolling 2D Standard Deviation method provide robust means to achieve acceptable segmentation results (IOU > 0.6, **Figure 3B**).

**Figure 3.**
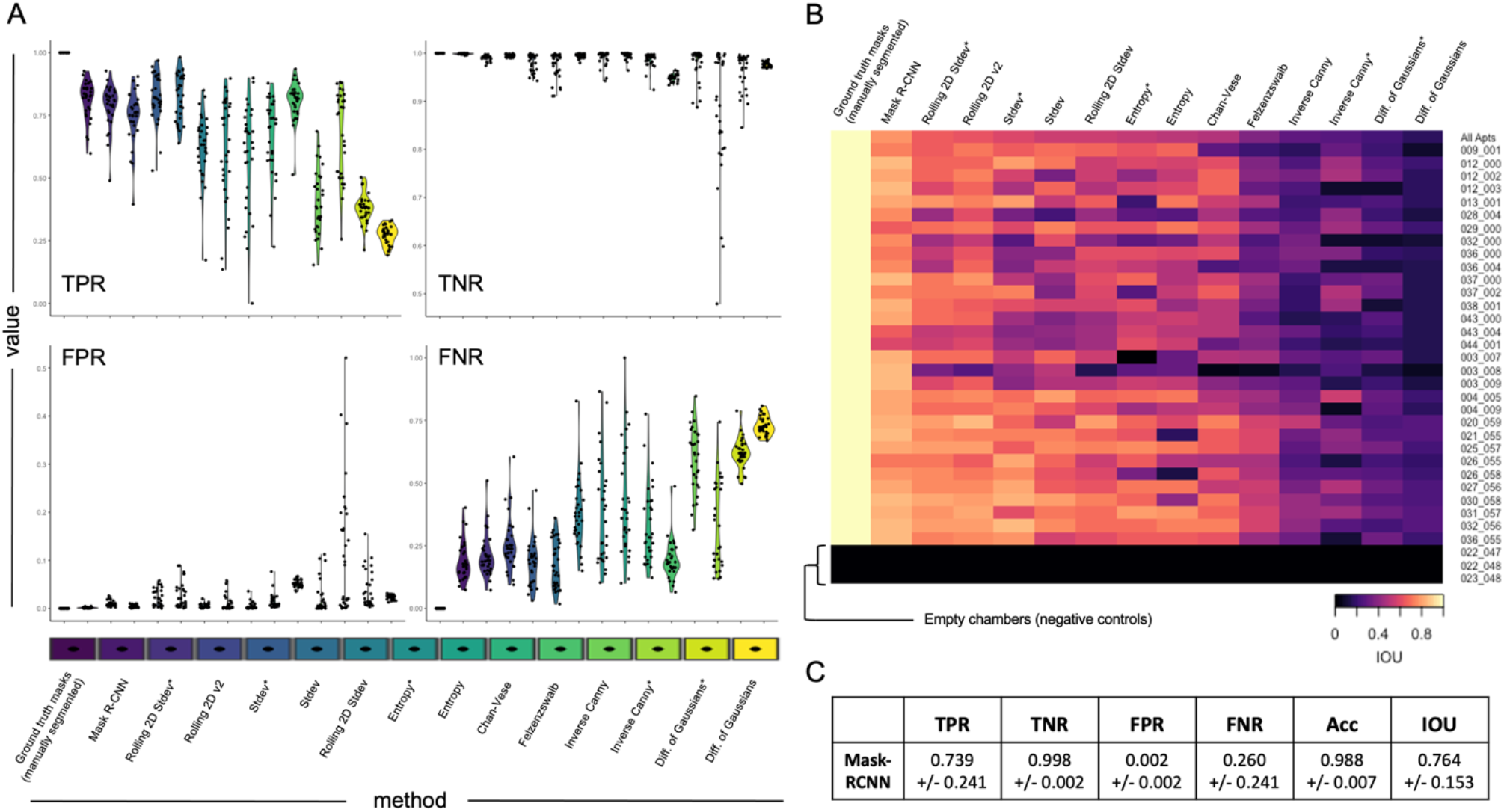
Method performance evaluation for test set images. (**A**) Pixelwise binary classification evaluation of cell segmentation masks form each method relative to manually segmented masks as ground truth. Plots show measured rates for each of 34 chamber images in the test set (dots) as well as the set-wide distributions (violins) for each method (TPR = True Positive Rate; TNR = True Negative Rate; FPR = False Positive Rate; FNR = False Negative Rate). (**B**) Calculated intersection over union (IOU) of test set cell segmentation masks and ground truth masks for each method ranked by overall performance. The last three rows represent negative control chambers that contained no cells. (**C**) Mean +/- standard deviation pixelwise classification rates relative to manual segmentation for the best overall method, mask-rcnn (Acc = Accuracy). Mask-rcnn overperforms the coarse-grained manual segmentation of cells in the ground truth masks, which contributes to an apparent reduction in TPR, Accuracy, and IOU.

Relative to reference classification, the Mask-RCNN model for cell detection achieves over 98% accuracy and IOU = 0.764 across test set images (**Figure 3C**). However, these values likely underestimate the method’s true performance, as the machine learning approach generates more precise and detailed segmentations of individual cells than captured in the manually generated bulk cell segmentation in the references. Beyond its superior classification performance, the machine learning approach has the significant advantage of yielding separate segmentation of each individual cell instance within an image rather than a single mask of cell-containing regions. This capability allows substantially greater cellular detail to be captured and quantified over the course of clonal growth experiments.

### Time-resolved clonal cell growth studies

Individual cells from an acute myeloid leukemia (AML) cell line (MOLM-13) were cultured on the Celldom™ array for 72 h under continuous media exchange conditions. Brightfield images were acquired periodically to visualize growth of individual clones within the population (**Figure 4A**). Time-resolved images reveal substantial variation in growth trajectories and clonal fates over time. By applying the mask-rcnn model for cell detection, clonal differences in proliferation (or death) are directly quantifiable from single cell counting. Notably, the model trained on instances of LCL 461 cells was capable of accurately detecting MOLM-13 cells with no additional training or adjustment. Chambers that each contain a single cell at the beginning of an experiment may exhibit highly variable cell densities at subsequent timepoints (**Figure 4B**). Over time, these differences manifest as per chamber cell distributions with increasing variance (**Figure 4C**), consistent with fates ranging from early cell death to maintenance of exponential growth by most or all clonal progeny.

**Figure 4.**
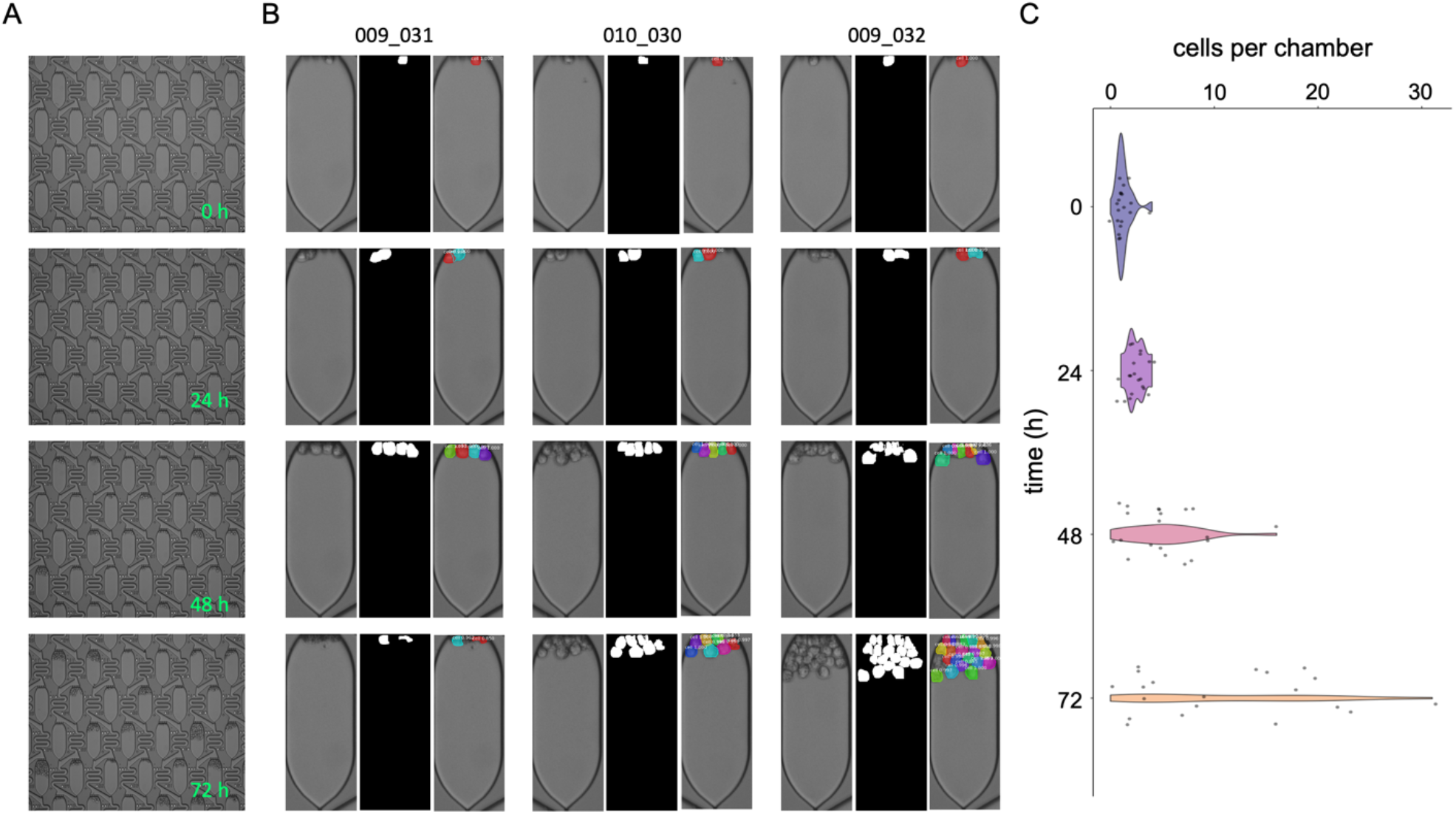
Time-resolved cell segmentation for chamber-indexed growth trajectories. (**A**) Example field of view depicting parallel single-cell culture outgrowth of MOLM-13 cells imaged at 24 h intervals over three days. The 0 h timepoint shows high-efficiency single-cell capture enabled by the microfluidic design. (**B**) Three selected chamber ROIs, binary masks, and individual cell instance from mask-rcnn segmentation of the time-resolved images presented in (A). The selected chambers demonstrate heterogeneity in clonal population growth rates and fates over time, as evidenced by the number of cells present in each chamber after 72 h. (**C**) Evolution of cell density in chambers over time. The number of cells in each chamber (dots) and distribution of cells per chamber (violins) in the given field of view are presented for each timepoint.

### Quantitative fluorescence imaging with single-cell resolution

The microfluidic culture array design is compatible with epifluorescence microscopy, which can be used to increase the dimensionality of clonal growth studies to include biomarker expression data (**Figure 5**). To demonstrate this capability, cultured MOLM-13 cells were stained on-chip for the pan-hematopoietic surface protein, CD45 (**Figure 5A**). Image co-registration across acquired channels enabled brightfield instance segmentations from the Mask-RCNN model to be used to quantify single-cell fluorescence intensity distributions in selected chambers (**Figure 5B**). Cell instance masks can be used in conjunction with simple or complex analyses of region properties to yield detailed morphological and statistical characterizations of single cell biomarker staining and to explore observed correlations (**Figure 5C**). This high-density phenotypic information can be compared within and across clones in culture chambers to interrogate aspects of diversity within cell lines of interest.

**Figure 5.**
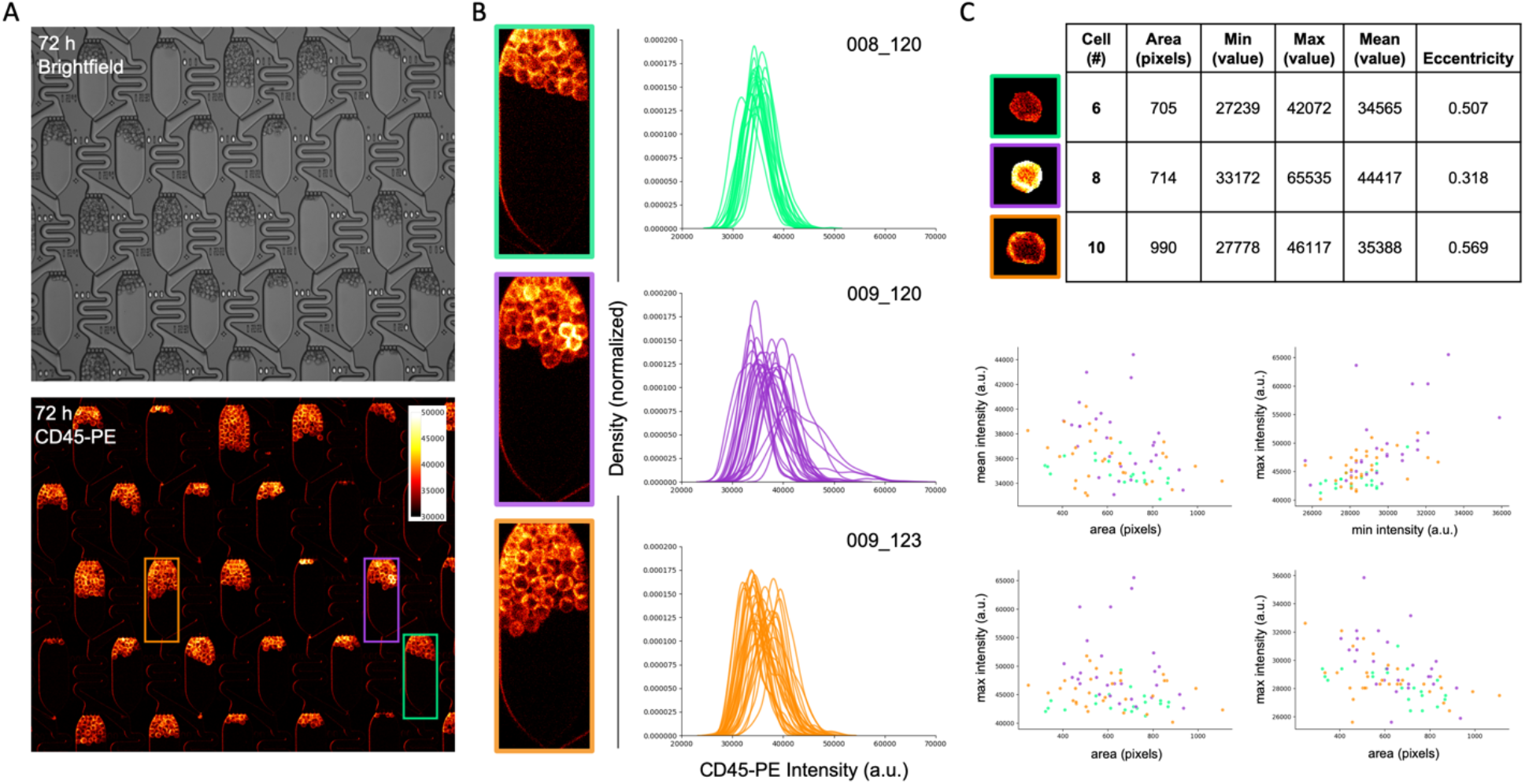
Clonal population morphology and biomarker expression characterization capabilities. (**A**) Brightfield (top) and fluorescence (bottom) imaging of MOLM-13 cells cultured on chip for 72 h. Live cell surface marker staining was performed at 72 h by spiking PE-conjugated anti-CD45 into the media at the microfluidic inlet. (**B**) Single-cell CD45-PE fluorescence characterization in individual culture chambers. Fluorescence intensity distributions of individual cells are plotted for each of three chambers of interest (green, purple, and orange ROIs). (**C**) Chamber-indexed deep phenotypic characterization through cell instance detection. High-accuracy single-cell segmentation enables extraction of individual cell sizes, shapes, and fluorescence intensity statistics (top), facilitating studies of intra- and inter-clonal phenotypic correlations (bottom).

## Discussion

Here we report a software package that extracts rich quantitation of individual cells imaged in a novel high-throughput clonal growth system. Importantly, we systematically evaluated numerous methods for cell segmentation and classification to determine the segmentation method that afforded the most accurate classification and highest information content for identified cells in the single-cell culture assay. Although several classical image processing techniques produced satisfactory classification results for cell-containing regions, a regional convolutional neural network was superior in key aspects of performance after minimal training.

Many of the tasks within the image processing workflow can be accomplished efficiently without machine learning models. For example, etched features of the chip (fiducial marks, addresses, chamber contours) are highly uniform and can be identified accurately using active contours or template matching since the target features are clearly defined *a priori*. By contrast, biological material is much more challenging to segment and quantify definitively, even for trained human analysts, in part due to its range of morphological presentations. Neural networks are favorable for the task of cell segmentation owing to both out-of-the-box performance as well as improvements and flexibility with respect to instance classification that can be achieved through further supervised model training. There have been numerous demonstrations of using machine learning for biological image analysis,^13,31-33^ however the application of such methods for analyzing high-throughput single-cell culture data is unique and opens a new frontier for innovative large-scale studies of single-cell biology. We expect that the initial cell detection model trained as part of this work can be adapted easily to accurately classify a broad variety of cells by end users on an experiment-specific basis. With sufficient training it may be possible to develop models that accurately distinguish different classes of cells by morphology, which would facilitate high-throughput investigations of interactions between distinct cell types in co-culture.

The ability to accurately count cells in highly parallel clonal cultures over time marks a significant advance to assay population-wide effects on growth of various stimuli and chemical perturbations. However, these methods can be leveraged for insights well beyond high-resolution dynamics of cell proliferation and death. Standard fluorescence imaging capabilities can be integrated to investigate functional correlations between individual clone dynamics and expression of proteins of interest. Additionally, we are investigating time-resolved gene expression studies on this platform using cell lines modified with fluorescent reporters and have recently developed protocols for on-chip fixing and permeabilization for end-point staining of intracellular targets (data not shown). Such studies offer an appealing synthesis of single-cell imaging and statistical power to evaluate heterogeneous expression and to gain deeper insight into phenotypic variance (and its implications) within nominally identical cell populations. Thus, we anticipate this combination of microfluidic platform and image analysis software is well-suited to diverse applications including compound screening, responses to stimuli, and fundamental behaviors and dynamics exhibited by thousands of individual cells in a population of interest.

## Materials and Methods

### Microfluidic Chips and Cell Culture

Microfluidic chips for single cell trapping and culture were designed and fabricated as described in Yellen *et al*, which also describes the protocol for loading and culturing cells in detail.^24^ Briefly, chips were assembled into microfluidic cassettes developed to enable imaging and continuous media exchange. After assembly, the microfluidic network was primed with 0.2 μm-filtered R10 media (RPMI + 10% fetal bovine serum (FBS), heat inactivated). Next, a single-cell suspension of the desired cells at 1×10^6^ cells/mL in filtered R10 was added to the microfluidic inlet and loaded deterministically into chamber traps. More media was added to rinse untrapped cells, and a brief pressure pulse was used to transfer cells from the traps into the chambers. After cell loading, chips were incubated with constant media exchange at 37°C and 5% CO_2_and imaged periodically.

### Cell Lines

Two cell lines were used in this study. A lymphoblastoid cell line (LCL 461) derived by transforming primary human B cells with Epstein-Barr Virus (EBV)^34-36^ was used to generate images for development and quantitative comparison of the segmentation methods, including mask-rcnn model training. An acute myeloid leukemia (AML) cell line with constitutively active FLT3 signaling (MOLM-13)^37^ was used for the time-resolved growth and fluorescence data presented in Figures 4 and 5.

### Live Cell Immunofluorescence

MOLM-13 cell clones cultured on-chip for 72 h were stained for the hematopoietic marker CD45 using a PE-conjugated primary antibody (mouse anti-human CD45-PE, Invitrogen #12-0451-82). Prior to imaging, 5 μL of labeled antibody was spiked into filtered R10 media at the chip inlet and flowed to stain the cells. Several volumes of media were then flowed to rinse residual unbound antibody, then brightfield and fluorescence images (Texas Red filter set) were acquired.

### Image Processing Code

The image processing workflow was implemented as a Python library that is freely available for use and modification (https://github.com/esorelle/cell-counter). The initial step of this pipeline uses OpenCV (cv2) to template match cross-shaped fiducial markers etched onto the microfluidic chip at precise locations. The centroid coordinates from each fiducial are used to construct sets of parallel lines for each image. The average slope of these lines is calculated and used to apply an image-wide rotation correction for small angles. Optionally, variation in light intensity across the image can then be normalized using a local adaptive background correction with a user-defined window size. Although this optional step provides remediation of non-uniform illumination, it reduces image contrast, which can adversely affect subsequent detection and segmentation steps if segmentation method hyperparameters are not adjusted accordingly. Following image corrections, rectangular regions of interest containing each single-cell culture chamber and its corresponding row and column address numbers are extracted based on their fixed positions relative to the fiducial mark centroids. Each numerical address ROI contains exactly three digits, which are parsed into three individual digit ROIs that are template-matched against a library of digit references to assign a consensus digit by majority vote. At this point in the pipeline, the correct address can be assigned to the extracted chamber and cell segmentation within the chamber ROI can be performed. The different cell segmentation methods developed and/or tested in this work are described by type below. Each method requires the extracted chamber ROI and an apartment reference mask (provided as a resource in the code library) as inputs.

#### Statistical Methods

Five segmentation methods based on simple image statistics were developed and tested. Generally, these methods were developed from the understanding that groups of pixels corresponding to cell features (e.g., internal morphology, cell membranes) have distinct spatial and intensity distributions relative to empty space within the extracted chamber image (i.e., background).

- Difference of Gaussians (DoG): this method is a pseudo difference of Gaussians. The method calculates a bilateral blur (which retains edge features) and a median blur (a less selective blur). Subtraction of the median blur image from the bilateral blur image preserves edges while removing noise. Blur images are cast to signed 16-bit images to accommodate negative values. The method requires two hyperparameters – median blur kernel size and bilateral blur kernel size. The bilateral blur kernel size should be less than or equal to the size of the feature of interest (single cell), and the median kernel size should be larger than or equal to the bilateral kernel. This method (and all others) also has a hyperparameter that sets a standard deviation threshold, which is set to 3.0 by default. We recommend that this threshold remain fixed for all methods.
- Standard Deviation (Stdev): this method is a standard deviation convolution implemented in MATLAB (https://www.mathworks.com/help/images/ref/stdfilt.html) adapted for Python. The method has one hyperparameter – the size of the convolution kernel.
- Entropy (Ent): this method is a straightforward implementation of the entropy calculation in scikit-image (https://scikit-image.org/docs/dev/auto_examples/filters/plot_entropy.html). The method has one tunable hyperparameter – the entropy kernel size.
- Inverse Canny Edge (ICE): this method is a simplified version of an inverse canny edge detection routine. First, a minimal blur (3×3 kernel) is applied, followed by a mild bilateral filter to enhance feature contrast. Next, a double threshold is applied to isolate regions above and below the value of a sample background region. The upper and lower threshold regions are combined to produce the segmentation mask. The method has one hyperparameter – the size of the median blur kernel.
- Rolling 2D Standard Deviation (r2d, r2d2): this method is like the Stdev method, except that it applied a rolling window function of the standard deviation separately along each row, then along each column of the extracted chamber image. The two resulting images are thresholded using a background sample, and the resulting masks are combined using a Boolean AND operation.

For each of these methods, the resulting cell segmentation mask is subsequently masked by the apartment reference mask (a rectangle of equivalent size to extracted chamber images that masks the contour of a typical apartment) to filter extraneous false positives outside of the culture chamber. The only exception is for version 2 of the Rolling 2D Standard Deviation method (r2d2), in which the reference mask filtering step is applied prior to the cell segmentation step.

#### Segmentation Algorithms

In addition to the five statistical methods described above, we tested scikit-image implementations of two established image segmentation routines – the Chan-Vese and Felzenszwalb algorithms. Briefly, the Chan-Vese algorithm^29^ seeks to find and segment features through active contours (https://scikit-image.org/docs/dev/api/skimage.segmentation.html#skimage.segmentation.chan_vese). The Chan-Vese algorithm requires tuning of two hyperparameters, lambda1 and lambda2. The Felzenszwalb algorithm^30^ is a graph-based method for image segmentation that oversegments images based on a scale parameter to generate fewer, larger segments or more, smaller segments (https://scikit-image.org/docs/dev/api/skimage.segmentation.html#skimage.segmentation.felzenszwalb). Once tuned for detection of cell-sized segments, larger and smaller segments (e.g., empty chamber area, noise, etc.) can be filtered out by size. As for the statistical methods, the resulting segmentation masks from these algorithms was filtered by an apartment reference mask to achieve the result for each chamber.

#### Machine Learning (regions with convolutional neural network)

The final method we tested was an adaptation of the popular Mask R-CNN package for object instance segmentation (https://github.com/matterport/Mask_RCNN).^31^ First, we used Visual Image Annotator^38^ (http://www.robots.ox.ac.uk/~vgg/software/via/via_demo.html) to generate a labeled single-cell training dataset from validation set images (n=70) containing more than 300 individual instances of LCL 461 cells. Next, we used the images and the corresponding annotation file (VIA-generate JSON) to train a model for single-cell detection. Interestingly, we were able to produce a reasonable cell detection model within 30 training epochs (100 steps per epoch) when initiated from a model previously optimized for the Common Objects in Context (COCO)^39^ image dataset (https://cocodataset.org/#home, ‘coco.h5’ file in the mask-rcnn library), despite the fact that the COCO dataset does not include any examples of grayscale, live cell microscopy images. Training was performed on a CPU with a 2.6 GHz processor (Intel Core i7) and 8 GB memory (run time: 56h 20m for 30 epochs). On the same machine, the time required to analyze a single image (chamber) varies from ∼30s to 90s depending on cell density.

### Segmentation Method Study Design

Individual chambers containing cells at various densities (including several empty chambers for negative controls) were extracted from five images using part I of the image processing workflow. These extracted chambers were divided into validation (n = 70 chambers) and test (n = 34 chambers) sets. Masks of cell-containing regions (not individual cells) were then generated for all validation and test images by manual segmentation for use as a ground truth reference. Except for the Mask R-CNN implementation, the hyperparameters for each method were optimized by iteratively evaluating performance in segmenting three of the validation images representing variable cell density. After hyperparameter tuning, masks for each method were generated for the validation set images and compared to reference segmentation masks to confirm performance. As described above, validation set images were used to generate single cell instance training examples for the Mask R-CNN model. At this point, no further changes were made to the methods, and each was applied to the test set images to generate the reported performance evaluation relative to segmentation reference masks.

### Method Performance Evaluation

Test set masks from each method were compared to reference segmentations to yield pixelwise measurements of classification performance: true positive rate (TPR, sensitivity, recall), true negative rate (TNR, specificity), false positive rate (FPR), false negative rate (FNR), accuracy (Acc), and intersection over union (IOU). Each of these values was calculated through Boolean operations implemented in Python. Individual image and set-wide classification results for each method were output as spreadsheets and imported into R for data visualization using the ggplot package. Mask-rcnn has the significant advantage of segmenting each individual cell which enables a deeper level of characterization, as shown in Figures 4 and 5. In the case of the mask-rcnn model implementation, composite instance detection masks were used for comparison against reference segmentation masks (and in most cases, outperform the reference segmentation). Individual instance masks were retained for single-cell morphology and biomarker expression profiles.

## Acknowledgments

We would like to thank Celldom, Inc. for access to prototype imaging, culture, and microfluidic systems that made this study possible. E.D.S. wishes to acknowledge funding from the Duke Viral Oncology Training Grant (NIH T32 # 5T32CA009111-42)

